# Paternal leakage of organelles can improve adaptation to changing environments

**DOI:** 10.1101/2020.12.18.423500

**Authors:** Arunas Radzvilavicius, Iain G. Johnston

## Abstract

Sexual eukaryotes have diverse mechanisms preventing the biparental inheritance of mitochondria and plastids, and reducing the coexistence of dissimilar organelle DNA (heteroplasmy). Nevertheless, paternal leakage often occurs in plants, fungi, protists and animals, and this leaves the possibility that heteroplasmy can in some contexts be advantageous. Theoretical models developed in the past revealed that maternal inheritance improves selection against deleterious mitochondrial mutations, but none of them have explained the observed variation in the extent of paternal leakage. Here we show that paternal leakage regulated by nuclear loci can evolve to maintain advantageous organelle diversity in fluctuating environments. Strict maternal inheritance reduces organelle variance within the cell, but this loss of diversity can be detrimental when environments are shifting rapidly. Our model reveals that high levels of paternal leakage can evolve in these types of rapidly changing environments and that strict maternal inheritance evolves only when the environment is changing slowly.

**Data:** Matlab/Octave implementation of the model is available at *Https://github.com/StochasticBiology/PaternalLeakageEvolution*.

## Introduction

Eukaryotes inherit their mitochondrial DNA (mtDNA) predominantly from one of the two fusing sex cells, the maternal gamete. Numerous examples illustrate the widespread nature of this maternal inheritance. In mammals, ubiquitin-mediated proteolysis and autophagy machinery removes sperm mitochondria not long after they enter the egg (Song et al., 2016). Fruit flies and other animals degrade their mtDNA in maturing sperm (Chan & Schon, 2012), fission yeast segregate maternal and paternal mitochondria into different meiotic spores formed from the zygote (Chacko et al., 2019), and bivalve mussels retain two sex-specific homoplasmic mitochondrial populations through differential segregation of mitochondria into somatic tissues and the germline (Cogswell et al., 2006; Ghiselli et al., 2011). In unicellular protists there are no morphological differences between maternal and paternal gametes, but they still have mechanisms preventing heteroplasmy, although far less is known about the specifics of the molecular machinery. In plants and algae, plastid DNA (ptDNA) is also inherited predominantly from one of the two fusing gametes (Birky, 2001; Bock, 2007).

The universal occurrence of uniparental inheritance (UPI) and other mechanisms selecting against heteroplasmy (Burgstaller et al., 2018) in all eukaryotic supergroups suggests that heteroplasmy has negative fitness effects and is therefore selected against (Christie et al., 2015). Recent experiments with mice showed that even low levels of heteroplasmy can impair cell function relative to the homoplasmic type (Sharpley et al., 2012; Latorre-Pellicer et al. 2019). In humans, heteroplasmy for some mtDNA mutations causes devastating mitochondrial disorders (Schwartz and Vissing, 2002; Wallace and Chalkia, 2013). Mathematical models revealed that maternal inheritance can evolve because it improves purifying selection by segregating out detrimental mitochondrial mutations (Hadjivasiliou et al., 2013; Radzvilavicius, 2016; Radzvilavicius et al., 2017). More efficient selection at the level of the cell can also protect mitochondrial DNA from Muller’s Ratchet (Muller, 1964; Radzvilavicius et al., 2017b), and it improves response to selection for rare advantageous mitochondrial mutations (Christie & Beekman, 2017). Although formulated with mitochondria in mind, these models are not limited to mtDNA transmission and can also explain the observed patterns of plastid DNA inheritance and their evolution.

Recent empirical evidence, however, demonstrates that uniparental inheritance is rarely absolute. Paternal leakage of mitochondria occurs in plants (Pearl et al., 2009; Woloszynska, 2010; McCauley 2013; Greiner et al., 2015), protists (Messenger et al., 2012), fungi (Xu and Li, 2015) and even in complex animals (Wolff et al., 2012, Nunes et al., 2013, Luo et al., 2018), and many plants inherit plastids from both parents (Birky, 2001; Bock, 2007). Overlooked by models based on purifying selection effects, the ubiquity of paternal leakage suggests an alternative hypothesis: rather than being solely the result of errors in mitochondrial elimination, low levels of heteroplasmy may confer some positive influence, and paternal leakage may be retained as the result of natural selection.

Maintenance of genetic diversity can be advantageous in populations to hedge bets against environmental change (Phillipi & Seger, 1989; Grafen, 1999), and provides a resolution to the tension between robustness and evolvability (Wagner, 2007). Because mitochondria are responsible for ATP production via oxidative phosphorylation, and OXPHOS enzymes are temperature-sensitive, mitochondrial haplotypes can perform differently depending on the prevailing environmental and thermal conditions (Dowling et al., 2007). It has been suggested that these gene-environment interactions may be responsible for the variation in mitochondrial genes among populations in distinct climatic conditions (Wolf et al., 2012), but the question of how sessile organisms with slow DNA evolution rates in their bioenergetic organelles can respond to environmental shifts remains unsolved. In this article we explore the hypothesis that organelle-environmental interactions may promote the maintenance of mtDNA and ptDNA polymorphism through selection for low levels of paternal leakage in shifting environmental conditions. We develop an evolutionary model, focusing on mitochondria but also applicable to plastids, in which the fitness of a cell depends on how well its mitochondrial haplotype matches the environment which fluctuates over time. We find that while UPI can evolve in constant and moderately changing environments, rapid fluctuations select for paternal leakage and biparental organelle inheritance.

### Model summary

Our modelling framework represents an evolving population of cells as matrices describing allele frequencies, and it uses linear algebra to model changes in mito-nuclear genotype distributions over time. Specifically, we use the transition-matrix formulation of mitochondrial-nuclear population dynamics first introduced by Radzvilavicius 2016. Here, we outline this approach, and present full details in Supplementary Methods.

To describe the allele frequencies in the evolving population at any time, we represent joint mito-nuclear genotypes in the two sexes as P-matrices ***P***^***f***^ (female subpopulation) and ***P***^***m***^ (male subpopulation), where an element 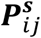 denotes the frequency of the joint mito-nuclear genotype with *i* mitochondria of haplotype *a*, and the nuclear allele *j* within the sex *s*. The dynamics of the system – genotype frequency changes – are described by the evolution of P-matrices as they are multiplied by transition matrices ***X***_***n***_: ***P***^***f***(***m***)^ (after) = ***X***_***n***_***P***^***f***(***m***)^(before). Each of the transition matrices represents an event that occurs in the organism’s lifecycle: the mutational matrix ***U*** element ***U***_*ij*_ represents the probability that a cell with *j* mutant mitochondria will have *i* mutants after the transition, and we construct similar matrices for selection, mitochondrial segregation at cell division and sexual reproduction (see Supplementary Methods). This framework provides exact descriptions of joint mitochondrial-nuclear genotype evolution between generations, and we seek solutions using standard numerical linear algebra methods and fixed-point iteration.

We model infinite populations of unicellular eukaryotes, each with *M* mitochondria and a haploid nucleus. Although here we focus on mitochondria, the same arguments can be applied to cells containing *M* plastids. A haploid nucleus models unicellular eukaryotes in which the diploid zygotic life cycle stage is transient, and the mating type or sex is determined at the haploid stage, represented by the unicellular alga *Chlamydomonas reinhardtii* and many yeasts, although the results also directly translate to diploids. Two mitochondrial haplotypes are segregating within the population, *a* and *A*, with the symmetric mutation rate between the two *μ*. Each of the two haplotypes is well matched to one of the two possible environments, *E*_*a*_ and *E*_*A*_, but performs poorly in the other.

Cell fitness depends on how well the overall mitochondrial population of the cell matches the current environment, *w*_env_ = 1 − (frequency of mismatched mitochondria)^*x*^. In environment *E*_*A*_, cell fitness is therefore 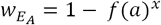 and in environment *E*_*a*_, the fitness of the same cell is 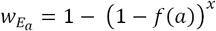, where *f*(*a*) is the frequency of *a*-type mitochondria within the cell. The nonlinearity *x* > 1 accounts for the mitochondrial threshold effects that result in increasing detrimental fitness effects of each additional mismatch mutation, consistent with empirical observations (Rossignol et al., 2003). The environment switches from *E*_*a*_ to *E*_*A*_ or *E*_*A*_ to *E*_*a*_ with a probability 1/*P* per generation, where *P* is the mean period of environmental fluctuations. Here *P* > 1, so that a given generation always experiences the same environment.

Each generation, the population goes through the life cycle of 1) mitochondrial mutation, 2) haploid selection, and 3) reproduction, which can be either clonal or sexual. Each of these stages is modelled by multiplying the matrix describing the current state of the system with a corresponding transition matrix. In sexual reproduction, cells of opposite mating types fuse at rate *r*_*sex*_. Because of the predominantly uniparental inheritance of mitochondria, all *M* mitochondria of the maternal mating type are transmitted to the zygote, while the paternal gamete transmits only *lM*, where paternal leakage *l* ≤ 1. Cell fusion is then followed by reductive cell division in which mitochondria are randomly distributed into two daughter cells. Cells can alternatively reproduce clonally (rate 1 − *r*_*sex*_), in which case the mitochondrial contents of the cell are duplicated and then distributed randomly across the two daughter cells.

The extent of uniparental inheritance is regulated by a haploid locus linked to the mating type (or sex-determination) locus. First, we determine the overall patterns of paternal leakage evolution, and then study how the levels of leakage and uniparental inheritance that evolve in model populations depend on environmental fluctuation and mitochondrial mutation rates.

## Results

In previous theoretical work we found that the overall pattern of paternal leakage evolution in populations with purifying selection depends on which sex regulates the extent of uniparental inheritance (Radzvilavicius et al., 2017a). We therefore first considered the evolution of paternal leakage via weak and rare mutations that increase or reduce paternal leakage by 1/*M* (see SI Methods). If the allele regulating paternal leakage is linked to the maternal mating type that, by definition, contributes more mitochondria to the zygote, we found that paternal leakage evolves either to the stable value of 0 (strict uniparental inheritance) or 1 (symmetric biparental inheritance, Fig. 1), depending on the initial condition. With paternal linkage of alleles regulating mitochondrial inheritance, on the other hand, intermediate levels of paternal leakage can evolve (Fig. 1). These differences arise because of the opposite ways in which an increase or reduction in the extent of paternal leakage affects the statistical linkage between the mitochondrial populations and the sex-linked loci (explained in detail by Radzvilavicius et al., 2017a). With maternal linkage of UPI regulation alleles, we studied how the frequency of UPI allele with *l* = 0 depends on mitochondrial mutation rates and environmental fluctuation frequency. With frequent environmental shifts (low *P*) and low mitochondrial evolution rates *μ*, biparental inheritance dominates (Fig. 2). Mitochondrial mixing associated with biparental inheritance results in weaker selection against mutations that are detrimental in one given environment, but it maintains within-cell polymorphism that improves adaptation when the environment changes unpredictably, that is, with low *P*. Uniparental inheritance, on the other hand, rapidly segregates out mitochondrial mutants maladaptive to the current environment, reducing within-cell diversity (Fig. 3). When there is a shift in environmental conditions, the lost haplotype that is now more fit must be regenerated through rare mutations, and this selects against uniparental inheritance in rapidly changing environments (Fig. 2).

**Figure 1.**
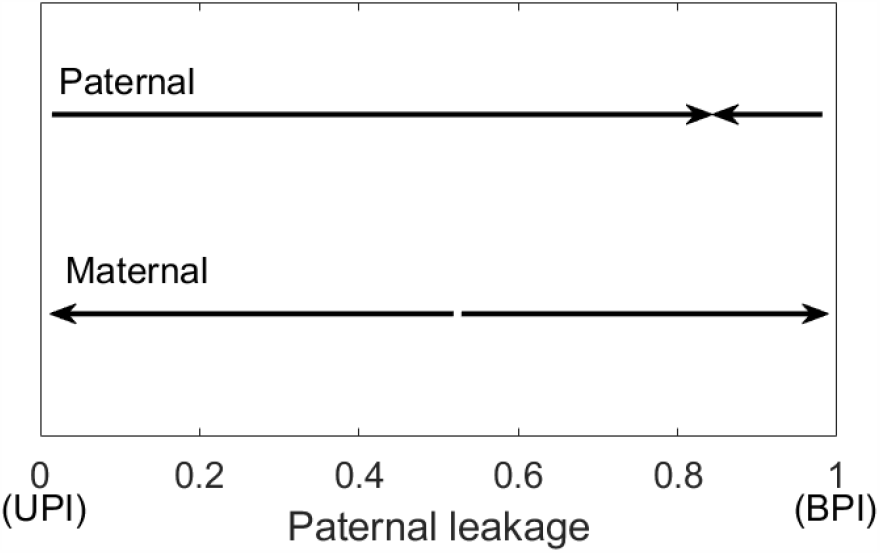
Direction of paternal leakage evolution via weak and rare mutations. In populations where mitochondria are inherited with paternal leakage l (horizontal axis) we tested if mutations that increase or reduce paternal leakage by 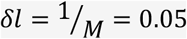 evolve, for all possible values of wild-type paternal leakage between 0 and 1. If the locus regulating the extent of paternal leakage segregates together with the maternal mating type (or female sex-determination) locus, only strict uniparental inheritance (paternal leakage of 0) or biparental inheritance (paternal leakage 1) can evolve. In taxa with paternal regulation of uniparental inheritance, paternal leakage evolves toward an intermediate value between 0 and 1. Sex rate is set to r_sex_ = 0.9, the mitochondrial mutation rates are μ = 0.01 (Maternal) and μ = 0.001 (Paternal), P = 100 and M = 20.

**Figure 2.**
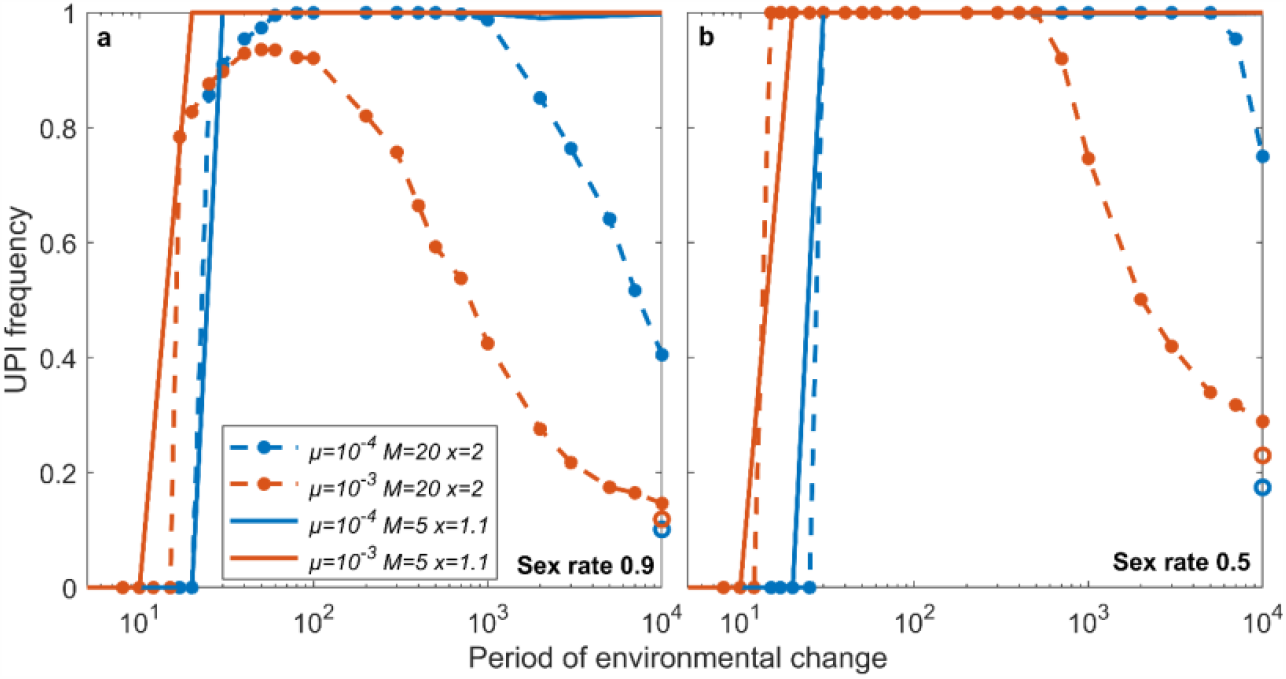
Mean frequency of the uniparental inheritance mutant allele (paternal leakage l = 0) linked to maternal mating type locus. In rapidly shifting environments and with low mitochondrial evolution rates, model populations evolve biparental mitochondrial inheritance as it preserves advantageous mitochondrial polymorphism in unpredictably changing environmental conditions. With completely uniparental inheritance, genotypes suited to a new environment would otherwise have to be regenerated through novel mutations. Uniparental inheritance evolves in slowly changing environments, where it improves response to selection by segregating out rare advantageous mutations. Finally, in constant environments (colored circles) the extent of UPI depends on the mitochondrial interaction nonlinearity x. With strong nonlinearities (x=2) heteroplasmy has the short-term advantage of masking detrimental fitness effects and dampens the evolution of UPI, while with weaker interactions (x=1.1) UPI increases mutational variance between cells and evolves to efficiently segregate out haplotypes that do not match the prevailing environment. In general, rare facultative sex (**b**) favors higher frequencies of UPI mutants relative to frequent sex (**a**), as it preserves a strong mitochondrial-nuclear linkage.

**Figure 3.**
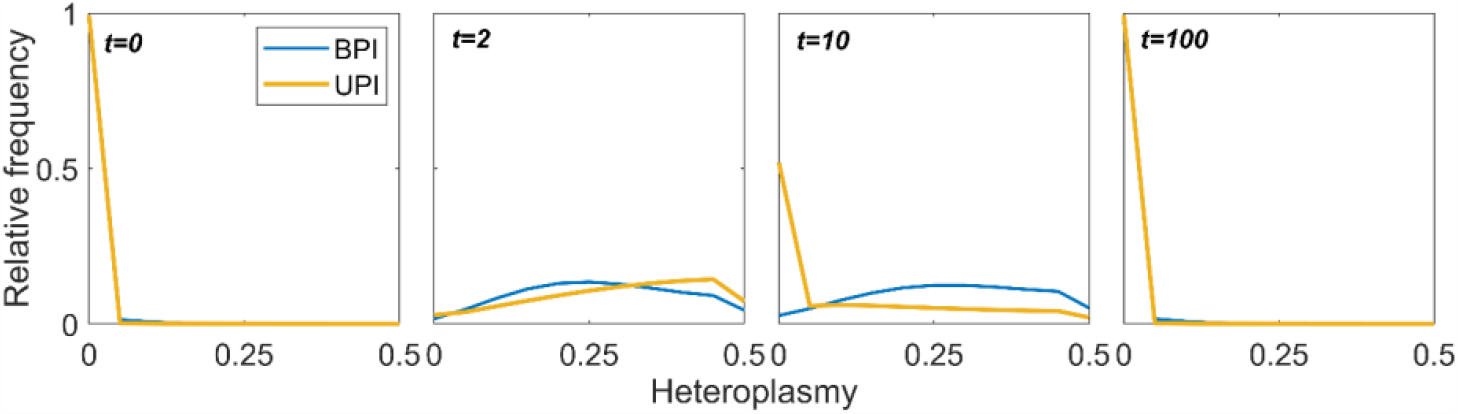
Mitochondrial heteroplasmy distributions at t=0, 2, 10 and 100 generations following the environmental shift. Heteroplasmy is maximized at 0.5 when both haplotypes are present within the cell at equal frequencies and it is 0 when only one haplotype – either a or A – is present within the cell. Cells with uniparental inheritance (yellow line) efficiently segregate advantageous and maladaptive mutations and experience a more rapid shift between haplotypes when there is an environmental change, while the biparentally reproducing subpopulation (blue line) lags behind and retains higher levels of heteroplasmy. With slow rates of environmental change, uniparental inheritance mutants will be more fit, but if the environmental changes are frequent (e.g. at t = 10), biparental inheritance will be advantageous as it retains the intracellular diversity that the uniparental mutants quickly lose. Mitochondrial mutation rate is μ = 0.0001, sex rate r_sex_ = 0.9. The frequency of UPI mutants is held constant at 10%.

With slower environmental changes, the increase in fitness associated with rapid segregation of rare advantageous mutations becomes more important than the retention of intracellular diversity, and the uniparental allele reaches high frequencies with low mutation rates. In constant environments, uniparental inheritance acts only to improve purifying selection by segregating out detrimental mutations, and so UPI alleles reach highest frequencies with rapid mitochondrial mutation accumulation and weakly nonlinear interactions between mitochondrial mutations *x*. In all cases, uniparental inheritance is most advantageous when sex is rare, because then linkage between mitochondrial and nuclear genomes is stronger (Fig. 2b).

Following our observation that paternal leakage can evolve to intermediate values with male-linked mitochondrial inheritance alleles, we next asked how the extent of paternal leakage that evolves through rare and incremental mutations depends on mutation rates and environmental shifts (Fig. 4). Once again, we find that when the environmental shifts are frequent, paternal leakage evolves to high values, as it maintains intracellular variance and improves adaptation when there is an environmental change (Fig. 4). However, in slowly changing environments, diversity can be regenerated through mutation and segregation, and so mitochondrial inheritance evolves towards lower values of paternal leakage, with rare sex and low mtDNA evolution rates selecting for more uniparental inheritance. Finally, the evolution of paternal leakage in constant environments is determined by purifying selection against detrimental mutations, and the short-term advantage of reduced intercellular variance that masks detrimental mutants (Radzvilavicius, 2016; Radzvilavicius et al., 2017a). Notably, these short-term effects are significant only with highly non-linear interactions between mitochondrial mutations (*x* = 2 in Fig. 4), and complete uniparental inheritance evolves in constant environments with weaker inter-mitochondrial interactions.

**Figure 4.**
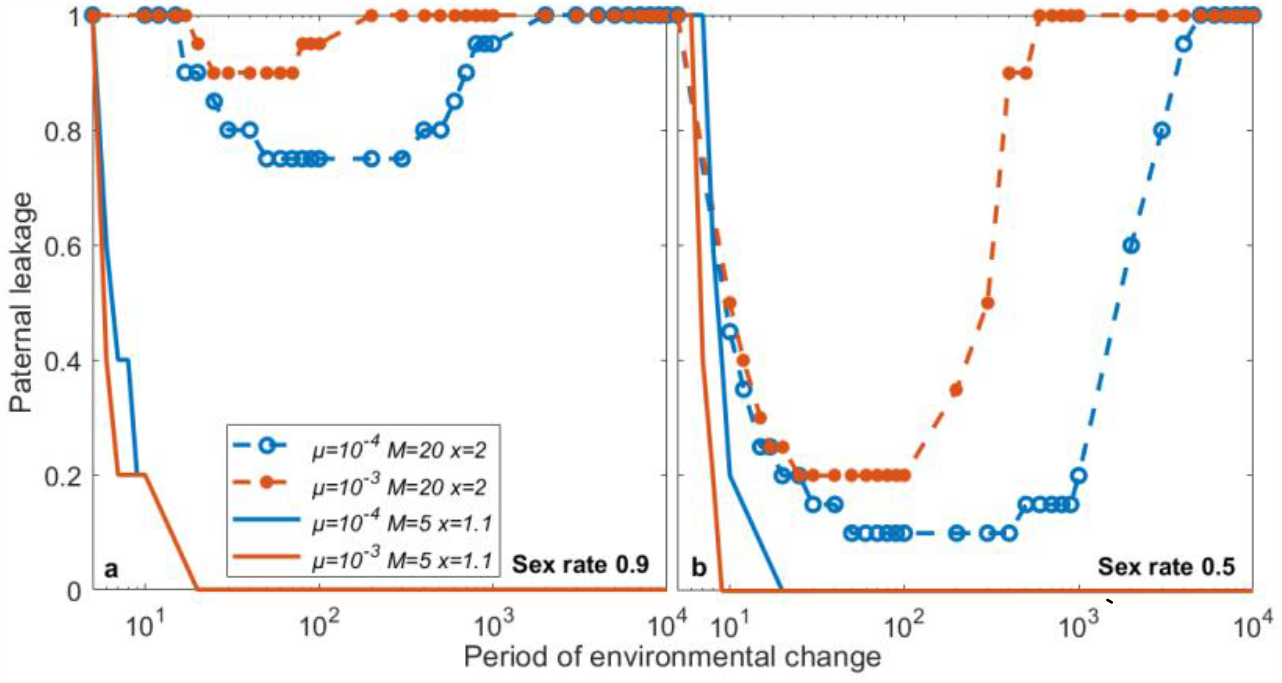
Evolution of paternal leakage in populations where uniparental inheritance is regulated by paternal alleles. High levels of paternal leakage evolve in rapidly changing environments, where, despite low variance across the population, mitochondrial haplotype polymorphism ensures rapid response to fast environmental shifts. Moderate rates of environmental fluctuations and low mtDNA evolution rates select for less paternal leakage, where adaptation to the shifting environments is driven by segregation of rare advantageous mutations. In constant environments, high levels of leakage evolve with strong interaction nonlinearities (x=2) because of the positive short-term effect of heteroplasmy masking detrimental fitness effects, and strict uniparental inheritance evolves with weaker nonlinear effects (x=1.1) because of its variance-increasing effect. Rare facultative sex (**b**) favors lower levels of paternal leakage relative to frequent sex (**a**), as it preserves a strong mitochondrial-nuclear linkage.

## Discussion

In recent years, the dogma of strict uniparental inheritance of mitochondria has been challenged by observations of persistent heteroplasmy in most eukaryotic supergroups, from protists to complex animals. We showed that if the performance of mitochondrial haplotypes depends on the external environment, then the extent of paternal leakage will depend on how frequent the environmental shifts are. In rapidly changing environments heteroplasmy provides within-cell diversity that is needed to rapidly adapt to the environmental change and paternal leakage evolves to maintain this advantageous polymorphism. However, because paternal leakage also reduces intercellular variance in heteroplasmy, it becomes detrimental in constant environments, where strict uniparental inheritance will evolve to efficiently segregate deleterious mutations. Selection against paternal leakage may be even stronger in natural populations because of the intrinsic negative fitness effects of heteroplasmy that are not related to the deleterious nature of mutations (Sharpley et al., 2012; Latorre-Pellicer et al, 2019).

Our theory of environmental selection for paternal leakage is supported by observations of more plastic modes of organelle inheritance and diverse organelle DNA structure in sessile organisms. Paternal leakage and mitochondrial genome recombination have been documented and inferred in plants (McCauley, 2013; Greiner et al., 2015), with strong evidence for biparental inheritance and leakage of plastids DNA (Birky, 2001; Bock, 2007), and there is an extraordinary diversity of mitochondrial inheritance modes and recombination rates in fungi (Yan et al., 2007; Wilson and Xu, 2012; Xu and Li, 2015). Sessile organisms are inevitably subject to environmental fluctuations and increased pressure on bioenergetic organelle genomes that interface with the environment (Johnston, 2019) and we believe that among these organisms there is a strong selection for mechanisms that rapidly respond to environmental fluctuations and regenerate advantageous mitochondrial and plastid diversity – paternal leakage, recombination, mitochondrial network formation and bottlenecking – even if constant environments would select against them.

These results have intriguing parallels with the evolvability and robustness argument that questions how genomes can escape negative mutational effects while maintaining flexibility to innovate (Wagner, 2007). When considering a population of organisms, the spread of genotypes through a genetic space can maintain the balance between robustness and evolvability. It is possible that multi-copy mitochondrial genomes, and mechanisms generating variance in haplotype distributions, may help resolve this paradox at the internal population level of organelles within an organism. The parallels extend into the evolutionary bet-hedging literature (Phillipi and Seger, 1989; Grafen, 1999), albeit at the intercellular, rather than the population level, where variance effects may be selected for in shifting environments even if constant environments would oppose their evolution.

The realization that paternal leakage may be a selected effect and not just an occasional breakdown of organelle removal machinery has profound implications for our understanding of eukaryotic genome evolution. Paternal leakage dampens or eliminates the evolution of detrimental sex-specific effects known as “mother’s curse” (Kuijper et al., 2015), and it may improve mitochondrial-nuclear coadaptation in the passive mating type. Frequent heteroplasmy may also affect population structure (McCauley, 2013) and reduce the accuracy of lineage history reconstructions that use mtDNA as a molecular marker.

In humans, heteroplasmy is associated with devastating mitochondrial disease phenotypes (Wallace and Chalkia, 2013) that appear when heteroplasmy reaches a certain threshold (Rossignol et al., 2003). These pathological heteroplasmies may be caused by paternal leakage (Luo et al., 2018), although the issue of paternal mitochondrial transmission in humans is surrounded by controversy (Salas et al., 2020; Wei et al., 2020). If paternal leakage is indeed regulated by unknown nuclear loci that experience selection, then mitochondrial disorders will be associated with the mutational variance at these loci. We therefore believe that a systematic search of nuclear regulators of paternal leakage across eukaryotes could improve and reshape our understanding of mtDNA disorders in the clinical setting and offer new therapeutic approaches. Critically, selection for and against paternal leakage may have significant but so far overlooked implications for the evolution of traits that affect intracellular mitochondrial diversity: eukaryotic life cycles and reproductive systems, the patterns of mitochondrial-nuclear coadaptation, sex rates, mating type numbers and germline structure in both sexes.

## Funding

This project has received funding from the European Research Council (ERC) under the European Union’s Horizon 2020 research and innovation programme (Grant agreement No. 805046 (EvoConBiO)).

## Notes

Https://github.com/StochasticBiology/PaternalLeakageEvolution

### Competing Interest Statement

The authors have declared no competing interest.

