## Supplementary Methods for "Paternal leakage of organelles can improve adaptation to changing environments"

**Supplementary Material**

### 1. Transition-matrix formalism of the joint mito-nuclear population dynamics

We model cells containing $M$ mitochondria of two possible haplotypes, $a$ and $A$. There are two sexes, male and female, and we model two nuclear alleles, wild-type ($j = 1$) and mutant ($j = 2$). In this work we represent a population of organisms with matrices $\boldsymbol{P}$, where matrix element $\boldsymbol{P}_{ij}$ denotes the frequency of the joint mito-nuclear genotype with $i$ mitochondria of haplotype $a$, $M - i$ mitochondria of haplotype $A$, and the nuclear allele of type $j$. We describe male and female organisms explicitly: $\boldsymbol{P^m}$ and $\boldsymbol{P^f}$ are the population matrices for male and female subpopulations.

The time evolution of $\boldsymbol{P}$ in infinite populations is then modelled using the transition matrix methodology introduced in Radzvilavicius, 2016. P-matrices are manipulated through multiplication by a set of square transition matrices $\boldsymbol{X_n}$, $\boldsymbol{P}(\text{next}) = \boldsymbol{X_n}\,\boldsymbol{P}(\text{current})$, where each of the transition matrices $\boldsymbol{X_n}$ is constructed to represent probabilistic life-cycle events: mutation, selection and reproduction. The explicit form of these matrices is described below. This framework gives exact changes in genotype frequency distributions between life cycle events and between generations (Radzvilavicius et al., 2017). In practice, we initialize all P-matrices with random mutational distributions and track their exact evolution using numerical linear algebra methods.

### 2. Mitochondrial mutation

Mitochondrial mutation between the haplotypes $a$ and $A$ is modelled as a binomial event with mutation rate $\mu$, which is symmetrical in both $a$-to-$A$ and $A$-to-$a$ transitions. The probability of $x$ new mutants in a cell with $j$ wild-type mitochondria then arises directly from the binomial distribution $\text{binom}(x, j, \mu) = \binom{j}{x}\mu^x(1-\mu)^{j-x}$, representing the probability of $x$ successes in $j$ trials with success probability $\mu$. With simultaneous bi-directional mutation between the two haplotypes, we combine two binomial terms to account for the possible different directions of change, and this combination gives rise to the form of the transition matrix $\boldsymbol{U}$. The probability that a cell will contain $i$ $a$-type mitochondria if it had $j$ $a$-type mitochondria before the mutational event is the matrix element $\boldsymbol{U}_{ij}$:

$$\boldsymbol{U}_{ij} = \sum_{x=0}^{j} \text{binom}(x, j, \mu)\text{binom}(x + i - j, M - j, \mu).$$

The first binomial represents the probability of $x$ $a$-to-$A$ mutations, and the second binomial samples $x+i-j$ $A$-to-$a$ mutations, so that the net increase in the number of $a$-type mitochondria is $i-j$. We then sum over all possible values of $x$. This mutational step of the life cycle transforms the P-matrices, $\boldsymbol{P^{s1}} = \boldsymbol{UP^{s}}$, where $s = m$ or $f$ and index 1 indicates the post-mutation life cycle stage within the given generation.

#### 3. Selection

There are two possible environments $E_a$ and $E_A$, and each is well-matched to either the mitochondrial haplotype $a$ or $A$ (as denoted by the subscript). Cell fitness depends on the total number of mismatches between the mitochondrial haplotype population within the cell and the current environment $n$: $w = 1-(n/M)^x$. We describe the fitness of different genotypes in a given environment as vectors: in environment $E_a$, fitness vector $\boldsymbol{W_a}$ element $\boldsymbol{W_{ai}}$ represents the fitness of the cell with $i$ $a$-type mitochondria ($M-i$ mismatches):

$$\boldsymbol{W_{ai}} = 1-\left(\frac{M-i}{M}\right)^x.$$

In environment $E_A$, fitness vector $\boldsymbol{W_A}$ has elements

$$\boldsymbol{W_{Ai}} = 1-\left(\frac{i}{M}\right)^x.$$

This nonlinear form of the fitness function with $x > 1$ models mitochondrial threshold effects, where the negative fitness effect of each new mismatch mutation increases with their total number, as fewer well-matched mitochondria are left to compensate, and it has been demonstrated experimentally (Rossignol et al., 2003).

The selection step then transforms P-matrices by resampling genotype frequencies weighted by their fitness, relative to the population mean in the current environment, $\boldsymbol{P_i^{s2}} = \boldsymbol{W}_{\mathbf{env}i}\,\boldsymbol{P_i^{s1}}/\langle \boldsymbol{W}_{\mathbf{env}}\rangle$, or in matrix form

$$\boldsymbol{P^{s2}} = \frac{diag(\boldsymbol{W}_{\mathbf{env}})\boldsymbol{P^{s1}}}{\sum_{ij} diag(\boldsymbol{W}_{\mathbf{env}})_{\boldsymbol{ii}}\,\boldsymbol{P_{ij}^{s1}}},$$

where the current environment env $= A$ or $a$, and $diag(\boldsymbol{X})$ is the square diagonal matrix with the entries of the vector $\boldsymbol{X}$ along the diagonal.

#### 4. Clonal reproduction.

Following selection, organisms undergo sexual reproduction at a rate $r_{sex}$, or otherwise reproduce clonally. In clonal reproduction (rate $1-r_{sex}$), cells double their mitochondrial populations from $M$

to $2M$, and then segregate them into the two daughter cells of equal size. This is described by sampling without replacement and follows the hypergeometric probability distribution. The hypergeometric distribution in general is given by

$$\text{hypergeom}(k, n, K, N) = \binom{n}{k}\binom{N-n}{K-k}\binom{N}{K}^{-1},$$

where $N$ is the population size, $K$ is sample size, $n$ is the total number of successes within $N$ and $k$ is the number of observed successes in $K$. If the parental cell before the clonal doubling step has $j$ mitochondria of haplotype $a$ (out of $M$ mitochondria in total) then the probability that a daughter cell has $i$ $a$-type mitochondria follows a hypergeometric distribution and the corresponding transition matrix $\boldsymbol{A}$ therefore has elements

$$\boldsymbol{A}_{ij} = \binom{2j}{i}\binom{2M-2j}{M-i}\binom{2M}{M}^{-1},$$

and the P-matrices after the asexual reproduction become $\boldsymbol{P}^{s3;asex} = \boldsymbol{A}\boldsymbol{P}^{s2}$, where sex $s = m$ or $f$, the index 3 indicates the life-cycle stage within the generation after reproduction, and the superscript *asex* indicates that the P-matrix was produced through asexual reproduction.

**5. Syngamy**

If the reproduction is sexual (rate $r_{sex}$), pairs of cells of opposite sexes fuse and mix their mitochondrial contents. The paternal gamete contributes $lM$ mitochondria to the zygote, sampled from the full haploid complement of $M$ without replacement. Again, we use the hypergeometric distribution to describe this sampling, giving rise to the transition matrix $\boldsymbol{H}^{\boldsymbol{l}}$ with elements

$$\boldsymbol{H}^{\boldsymbol{l}}_{ij} = \binom{j}{i}\binom{M-j}{lM-i}\binom{M}{lM}^{-1}$$

that represent probabilities that the male gamete with $j$ $a$-type mitochondria will contribute $i$ $a$-type mitochondria to the zygote. The maternal gamete contributes all of its $M$ mitochondria to the zygote, which after syngamy contains $lM + M$ mitochondria. To restore the full diploid complement of $2M$, we sample again, this time with replacement, which necessitates using the binomial distribution $\text{binom}(i, K, p) = \binom{K}{i} p^x (1-p)^{K-i}$, defining the probability of $i$ successes in $K$ samples with success probability $p$. We therefore obtain the matrix $\boldsymbol{B}^{\boldsymbol{l}}$ of binomial transition probabilities:

$$\boldsymbol{B}^{\boldsymbol{l}}_{ij} = \binom{2M}{i}\frac{j}{lM+M}^{i}\left(1-\frac{j}{lM+M}\right)^{2M-i}.$$

After sexual cell fusion, and before the binomial resampling, the number of $a$-type mitochondria in the zygote $i$ represents all possible combinations of mutant numbers in the two gametes that add up to $i$:

$$\boldsymbol{K}_i = \sum_j \boldsymbol{k}_j^{\boldsymbol{m}} \boldsymbol{k}_{i-j}^{\boldsymbol{f}},$$

where $\boldsymbol{k}^{\boldsymbol{m}}$ and $\boldsymbol{k}^{\boldsymbol{f}}$ are vectors representing resampled male and female gametic contributions with a desired nuclear allele, and their elements $\boldsymbol{k}_i^{\boldsymbol{f}(\boldsymbol{m})}$ denote frequencies of gametes with $i$ $a$-type mitochondria out of $M$ (female gamete) or $lM$ (male gamete). The expression defines vector convolution, $\boldsymbol{K} = \boldsymbol{k}^{\boldsymbol{m}} * \boldsymbol{k}^{\boldsymbol{f}}$. The joint mito-nuclear genotype distribution in diploid zygotes of the wild-type (no mutant nuclear alleles) are then

$$\boldsymbol{Z}^{\boldsymbol{l}} = 2\boldsymbol{B}^{\boldsymbol{l}} \left(\boldsymbol{H}^{\boldsymbol{l}} \boldsymbol{P}_{:\mathbf{1}}^{\boldsymbol{m}\mathbf{2}} * \boldsymbol{P}_{:\mathbf{1}}^{\boldsymbol{f}\mathbf{2}}\right),$$

where the index notation $:i$ here and later represents the $i$-th column of the given matrix.

If paternal leakage is regulated by maternal nuclear alleles, the genotype distributions in the mutant zygotes (a single mutant nuclear allele linked to either male or female sex determination) are $\boldsymbol{Z}^{\boldsymbol{L}} = 2\boldsymbol{B}^{\boldsymbol{L}} \left(\boldsymbol{H}^{\boldsymbol{L}} \boldsymbol{P}_{:\mathbf{1}}^{\boldsymbol{m}\mathbf{2}} * \boldsymbol{P}_{:\mathbf{2}}^{\boldsymbol{f}\mathbf{2}}\right)$, and if it is regulated by nuclear loci linked to male sex determination, $\boldsymbol{Z}^{\boldsymbol{L}} = 2\boldsymbol{B}^{\boldsymbol{L}} \left(\boldsymbol{H}^{\boldsymbol{L}} \boldsymbol{P}_{:\mathbf{2}}^{\boldsymbol{m}\mathbf{2}} * \boldsymbol{P}_{:\mathbf{1}}^{\boldsymbol{f}\mathbf{2}}\right)$, where $L$ is paternal leakage in the mutant subpopulation.

**6. Meiosis**

Diploid zygotes with $2M$ mitochondria undergo two meiotic divisions back into the haploid phase with $M$ mitochondria. In the first subdivision, the mitochondrial populations are doubled and then segregated into the two daughter cells. This is modelled as sampling without replacement following hypergeometric probability density as before, and the corresponding transition matrix $\boldsymbol{L}^{\mathbf{1}}$ then has elements

$$\boldsymbol{L}_{ij}^{\mathbf{1}} = \binom{2j}{i}\binom{4M-2j}{2M-i}\binom{4M}{2M}^{-1},$$

that once again denote probabilities that a zygote with $j$ $a$-type mitochondria will have $i$ $a$-type mitochondria after the transition. The second meiotic subdivision reduces the number of mitochondria back to $M$, again through hypergeometric sampling:

$$\boldsymbol{L}_{ij}^{\mathbf{2}} = \binom{j}{i}\binom{2M-j}{M-i}\binom{2M}{M}^{-1}.$$

The P-matrices after the sexual reproduction then become

$$\boldsymbol{P}_{:\mathbf{1}}^{\boldsymbol{s}\mathbf{3};\boldsymbol{sex}} = \boldsymbol{L}^{\mathbf{2}}\boldsymbol{L}^{\mathbf{1}}(\boldsymbol{Z}^{\boldsymbol{l}} + \boldsymbol{Z}^{\boldsymbol{L}})$$

for both sexes $s$, and

$$\boldsymbol{P}_{:\mathbf{2}}^{\boldsymbol{sr}\mathbf{3};\boldsymbol{sex}} = \boldsymbol{L}^{\mathbf{2}}\boldsymbol{L}^{\mathbf{1}}\boldsymbol{Z}^{\boldsymbol{L}}$$

for the sex $sr$ (male or female) that regulates paternal leakage through sex-linked nuclear alleles.

At the start of the next generation the mito-nuclear genotype frequencies are the sum of sexual and asexual contributions,

$$\boldsymbol{P}^{s} = r_{sex}\,\boldsymbol{P}^{s3;sex} + (1 - r_{sex})\boldsymbol{P}^{s3;asex}.$$

**7. General patterns of paternal leakage evolution**

To produce the main text Figure 1, for each possible value of wild-type paternal leakage $l$ we studied the evolution of paternal leakage mutants $L = l \pm 1/M$ (where $M$ is the total number of mitochondria in the cell, and the smallest difference in paternal leakage corresponds to a single paternal mitochondrion). Because these are mutations with weak fitness effects, their fitness advantage is approximately independent of their frequency. Therefore, they either uniformly increase in frequency and replace the wild-type allele, or their frequency decreases towards zero. The long-term evolutionary patterns of paternal leakage can therefore be described through rare and weak mutations, each reaching the frequency of 1 or 0 before the next mutation arises (Radzvilavicius et al., 2017).

For each pair of $l$ (wild-type paternal leakage) and $L$ (mutant paternal leakage) we first solved the life-cycle system of matrix equations for the steady-state P-matrices at mutation-selection equilibrium with only the wild-type allele $l$, so that $\boldsymbol{P}^{s}_{:2} = \boldsymbol{0}$ for both sexes. This equilibrium is defined by the self-consistency requirement, where at generation $k$, $\boldsymbol{P}^{s}(k-1) = \boldsymbol{P}^{s}(k)$, and it is independent of initial conditions. We solve for self-consistency using fixed-point iteration with random initial conditions with the numerical accuracy of $10^{-10}$. This means that genotype frequencies stay constant between generations within the range of $10^{-10}$.

In constant environments, self-consistency is achieved in ~60 generations for $\mu = 0.1$ , $l = 0$, $r_{sex} = 0.9$, $M = 20$, and in ~240 generations for $\mu = 0.001$, $l = 0.1$, $r_{sex} = 0.9$, $M = 20$. In changing environments the P-matrices may not reach an equilibrium between the successive environmental shifts, and then we track the evolution of P-matrices for $k = 5000$ generations until they converge to representative behaviour, which is at least 10-times the time needed to converge to equilibrium in constant environments for our parameter values.

Then, we model mutations in paternal leakage regulation that arise at frequency $f$ in generation $k + 1$:

$$\boldsymbol{P}^{sr}(k+1) = \left(\begin{bmatrix} 1-f & 0 \\ f & 1 \end{bmatrix} \boldsymbol{P}^{sr}(k)^{T}\right)^{T},$$

so that the mutant distribution in the sex regulating paternal leakage is $\boldsymbol{P}^{sr}_{:2}(k+1) = f\boldsymbol{P}^{sr}_{:1}(k)$ and $\boldsymbol{P}^{sr}_{:1}(k+1) = (1-f)\boldsymbol{P}^{sr}_{:1}(k)$.

Although the fitness advantage of the new mutants is frequency-independent, we use the initial frequency of $f = 0.01$, and then track the fate of the mutant as it evolves to the frequency of 1 (replacing the wild-type allele) or the frequency of 0 (Fig. S1). Main text Figure 1 represents the summary of these calculations for all pairs of $l$ and $L$, where the direction of an arrow represents mutant replacement and the opposite direction represents stability against new mutations in paternal leakage. Main text Figure 3 reports full distributions $\boldsymbol{P^f}$ as they evolve after an environmental shift, while keeping the frequency of $L$ mutants at a constant value of 20%.

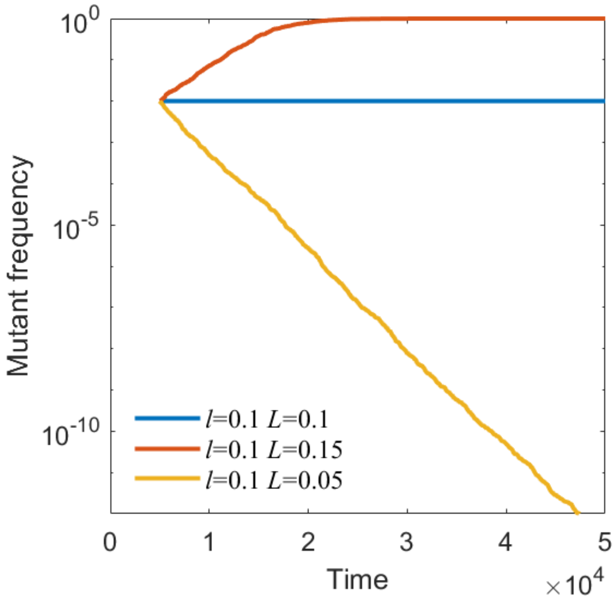


**Figure S1**. Evolution of paternal leakage mutant $L$ in the wild-type population with paternal leakage $l$. $P = 100$, $\mu = 0.001$, paternal, $M = 20$, $r_{sex} = 0.9$. The mutant either replaces the wild type allele or evolves to the frequency of 0. This analysis represents a single point in main text Figure 1, where it is repeated for all values of $l$ between 0 and 1.

### 8. Evolution of uniparental inheritance with maternal regulation

Our analysis, summarized in main text Fig. 1, showed that when paternal leakage $l$ is regulated by a nuclear locus linked to female sex-determination locus, only $l = 0$ (strict maternal inheritance) or $l = 1$ (symmetric biparental inheritance) can evolve (and are stable against weak $l$ mutants). W therefore limited our analysis with maternal regulation to $L = 0$ mutant evolving in the wild-type population of $l = 1$. The fitness advantage of this mutant is now frequency-dependent, and therefore a stable polymorphism is possible.

As before, an $L = 0$ mutant arises at a frequency $f = 0.01$ after $k = 5000$ generations. We then track the evolution of P-matrices over 5000 environmental fluctuation periods, which our manual examination shows is long enough converge to representative behaviour, and then we calculate the mean mutant frequencies over half of its history length (at least 2500 environmental fluctuation periods). These calculations for different parameter values are summarized in main text Figure 2.

### 9. Evolution of paternal leakage with male-linked nuclear regulation

As we demonstrate in main text Figure 1, when there is paternal regulation of mitochondrial inheritance, intermediate values of paternal leakage can evolve (and are then stable against rare mutants in the neighborhood of $l$). For evolution of paternal leakage with male-linked regulation alleles, we expanded our analysis to all possible pairs of $l$ (between 0 and 1) and $L = l \pm 1/M$. We repeated our analysis of SI Section 7 for different parameter values, looking for a value of $l$ such that both mutants $L = l - 1/M$ and $L = l + 1/M$ do not spread, but $l$ itself evolves from the close values of $L$ (Fig. S2). The summary of this analysis is reported in main text Figure 4.

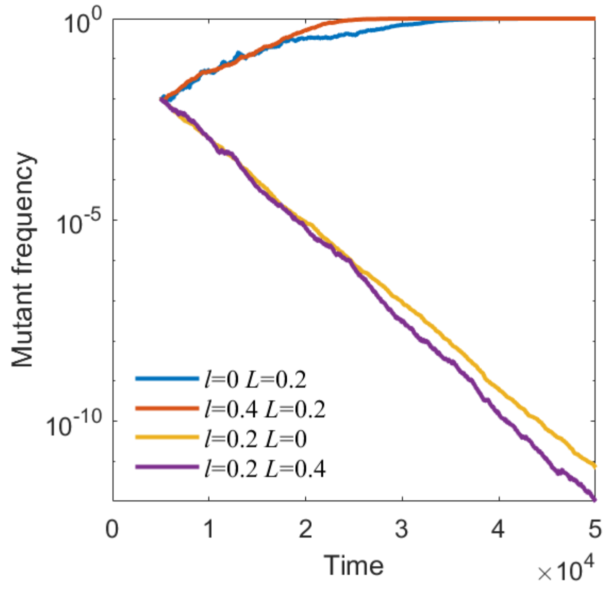


**Figure S2**. Evolution of paternal leakage mutant $L$ in the wild-type population with paternal leakage $l$, illustrating the stability of $l = 0.2$. $P = 10$, $\mu = 0.001$, $M = 5$, $r_{sex} = 0.9$. This analysis represents a single point in main text Figure 4, where it is repeated for all values of l between 0 and 1 and different parameter values to determine stable values of paternal leakage.
